# Sleep EEG distinguishes learned from novel auditory cues without predicting memory retention in a targeted memory reactivation study

**DOI:** 10.64898/2026.11.13.766368

**Authors:** Liisa Raud, Martine Moen, Markus Sneve, Solvår Jørgensen, Øystein Sørensen, Emilie Sogn Falch, Björn Rasch, Anders Fjell

## Abstract

**Study Objectives:** Memory consolidation during sleep is thought to build on processes initiated during encoding, but how encoding-related brain activity affects individual memories during sleep is unknown. Using targeted memory reactivation (TMR), we tested whether auditory cues elicit memory-related responses during sleep, whether encoding theta and delta power predict these responses, and whether encoding- and sleep-related measures predict long-term retention.

**Methods:** Eighty-one healthy young adults learned sound-associated item–face/place pairs during electroencephalography (EEG). Memory was tested before sleep, 12 hours, and 5 days after encoding. During non-rapid eye movement sleep, a subset of learned sounds and novel control sounds were replayed. Item-level mixed-effects models assessed behavioural and EEG predictors of retention success, and item-level correlations related encoding to cue-induced power during sleep.

**Results:** Pre-sleep memory strength and encoding vividness predicted retention, with the vividness effect depending on encoding theta power. TMR did not improve memory and was associated with poorer retention of initially weak memories. Learned cues elicited stronger delta responses than novel sounds during sleep, yet cue-induced delta, theta, and sigma power did not predict retention. Encoding and sleep EEG power were unrelated and showed distinct scalp topographies.

**Conclusions:** In a large item-level dataset, the sleeping brain distinguished learned from novel cues, but this processing did not translate into durable retention, which was instead determined at encoding. Encoding-related oscillations did not relate to memories preferentially reactivated or strengthened by sleep cueing. These findings caution against treating cue-induced sleep oscillations as markers of successful reactivation or consolidation.

## Introduction

Most everyday memories fade rapidly, whereas a subset become durable over days and weeks. A mechanistic account of this selectivity must explain how neural processes at encoding interact with offline consolidation to determine which traces are stabilized. Classical theories and supporting empirical work suggest that memories are formed via hippocampal–cortical communication during encoding^1,2^, and consolidated via subsequent spontaneous reactivation of these traces, particularly during non-rapid-eye-movement (NREM) sleep^3,4^. Both encoding and consolidation processes express rhythmic, oscillatory brain activity that, to some extent, can be indexed via scalp electroencephalography (EEG). However, no direct link has been established between oscillatory rhythms during encoding and sleep.

During encoding, theta oscillations (4–8 Hz) are thought to coordinate hippocampal–cortical communication by organizing neural activity into recurring windows of high excitability, during which synaptic potentiation is preferentially induced^5^. Although theta is most prominent in the hippocampus^6–9^, the fronto-medial theta oscillations measured by scalp EEG synchronize with hippocampal theta during memory and navigation tasks^10,11^. In addition, fronto-medial theta also arises during other cognitive tasks, putatively indexing domain-general control processes^12,13^, and is therefore best understood as a cortical signal that only partially reflects hippocampal–prefrontal interactions. Increases in scalp-recorded theta power are thought to reflect synchronous activity in larger neural populations and synchronized output is more effective at driving downstream targets^14,15^, so fronto-medial theta power may index how strongly a region influences the wider memory network. In line with this, theta power during encoding predicts subsequent remembering^16^, scales with confidence^17^ and future relevance^18^ of memories, and may tag memories for prioritized consolidation^19,20^. Delta activity (1–4 Hz), linked to attentional and contextual control during wakefulness^21^, is a further candidate for binding information across sleep and wakefulness. In NREM sleep, slow oscillations in the same frequency range have been linked to consolidation, and their regional power increases during sleep over cortical areas engaged by prior learning^22^. Encoding-related theta or delta power may therefore indicate the activation of the cortical networks later recruited for memory consolidation during sleep.

According to active systems consolidation theory, newly encoded hippocampus-dependent memories are repeatedly reactivated during both quiet wakefulness and sleep, gradually becoming integrated into long-term neocortical networks^3,4^. NREM sleep appears particularly favourable for this transformation, during which cortical slow oscillations, thalamocortical sleep spindles, and hippocampal sharp-wave ripples jointly create temporally precise windows for replay and plasticity^23–25^. Targeted memory reactivation (TMR) provides a powerful tool to probe this mechanism by re-presenting sensory cues linked to prior learning during NREM sleep, thereby biasing which memories are reactivated^26^. TMR generally improves subsequent memory for cued items^27^ with effects persisting over several days^28–31^, in line with a causal role of sleep reactivation in consolidation. Auditory cues are particularly well suited to this approach, as the sleeping brain continues to evaluate incoming sounds for their relevance^32,33^. Accordingly, sounds linked to prior learning elicit stronger responses than unlearned control sounds, including increased parahippocampal activity^34^, enhanced theta and spindle activity^35,36^, and late item-specific EEG representations that predict later memory^37^.

Within this framework, theta- or delta-band activity may act as candidate markers bridging wake encoding and sleep. Although the theoretical model emphasizes slow oscillations, spindles, and ripples, TMR studies additionally report increases in theta power during sleep in response to memory cues^38,39^. The presence of comparable theta and delta power changes during both wakeful encoding and sleep may indicate a common, state-independent signature of memory-related network activity. Whereas the sleep cueing benefit seems to be modulated by prior encoding strength^40,41^, links between encoding and sleep physiology have so far been reported mainly for supposed spontaneous consolidation during sleep: theta power during learning predicts subsequent spindle density^19,42^ and slow oscillation–spindle coupling^43^. Whether encoding-related power also predicts the neural response during the reactivation of the same memories during sleep remains unknown.

Here, we conducted an overnight auditory TMR study, expecting a positive effect of memory retention, and further addressing three questions. First, we asked whether auditory cueing during NREM sleep elicits memory-related neural responses, contrasting cues linked to prior learning with new control sounds. Second, we asked whether the strength of these cue-evoked responses is associated with oscillatory power during associative encoding. We contrasted two competing scenarios. If synchronized cortical engagement during encoding tags memories for later consolidation, items encoded with higher theta and delta power should elicit stronger cue-evoked theta, delta, or spindle-band (sigma) power during sleep, yielding a positive relationship between encoding and sleep power. Alternatively, if sleep-dependent consolidation compensates for weak encoding, TMR should preferentially benefit weakly encoded items, which would elicit stronger cue-evoked responses despite lower encoding power, yielding a negative relationship. Lastly, we asked whether memory retention is predicted by encoding-related factors (i.e. neural response, subjective vividness, and initial encoding strength), by the neural response to the auditory cued during sleep, or by a combination of those. Critically, we tracked each item individually, relating its retention 12 hours and 5 days later to encoding and sleep measures.

## Methods

### Sample

Out of 90 participants recruited for the study, the analysis was performed on 81 participants (60 females, 21 males, age range 21-40, mean age = 30, SD = 4.7). The 9 participants were discarded due to inability of falling asleep or frequent awakenings due to the TMR procedure. All participants were right-handed, reported no sleep disturbances over the last six weeks, reported routine sleep schedules, were not diagnosed with any psychiatric or neurological conditions, and had not been taking any medications that affect the nervous system in the last year. Participants were asked to not drink caffeinated beverages before the experiment and to avoid naps during the daytime. Compliance was checked upon the arrival to the laboratory, and it was noted that most, though not all, participants complied with these instructions.

Before recruitment, we conducted power analyses at both the group and item levels in R (version 3.6.0) using the *simr*^44^ package. At the group level, a TMR benefit of Cohen’s d = 0.25 was assumed, based on a meta-analytic estimate for within-participant designs using auditory verbal cues^27^. A paired t-test indicated that approximately 80 participants would provide 80% power at α = 0.05. However, as item-level analysis was planned, we simulated trial-wise memory accuracy using mixed-effects logistic regression, with cueing condition as the predictor of interest, pre-sleep alternative forced choice (AFC-1) accuracy as a covariate, and participant-specific random intercepts. Parameters were derived from the young sleep-condition sample of a previous study using the same memory paradigm but without TMR cues (n = 45)^45^. We specified an odds ratio of 1.26, corresponding to a 0.25 SD improvement in the baseline accuracy. Assuming 64 cued and 64 uncued items per participant, n = 46 provided ≈80% power. Data collection continued until n = 90 because the protocol included a simultaneous EEG–fMRI follow-up session (not reported here), for which additional recruitment was required to accommodate attrition.

### General procedure

Participants arrived at the lab at approximately at 20.30 in the evening (Figure 1A), though the exact timing was adjusted for each participant to accommodate their habitual sleep times. They were fitted with EEG caps and performed a memory encoding task, followed by an immediate forced choice learning test (8AFC-1). They were then allowed a break for getting ready to sleep and directed to a different room for sleeping. The EEG electrodes were re-checked, and the research staff stayed in an adjacent room, monitoring the EEG signal throughout the night. Participants went to bed at around 23.00. If stable NREM sleep stage 2 or 3 was detected, by manual monitoring of the data, the researchers started the TMR procedure. Upon the completion of all 12 TMR cycles, participants were allowed to sleep until 6.00 in the morning. After awakening, a small breakfast was offered, and participants then completed two memory tests: a category level retrieval test and an exemplar-level forced choice task (8AFC-2). This marked the end of the experimental session. After five days, participants received an online version of the exemplar-level forced choice task (8AFC-3), which was completed at their convenience.

**Figure 1.**
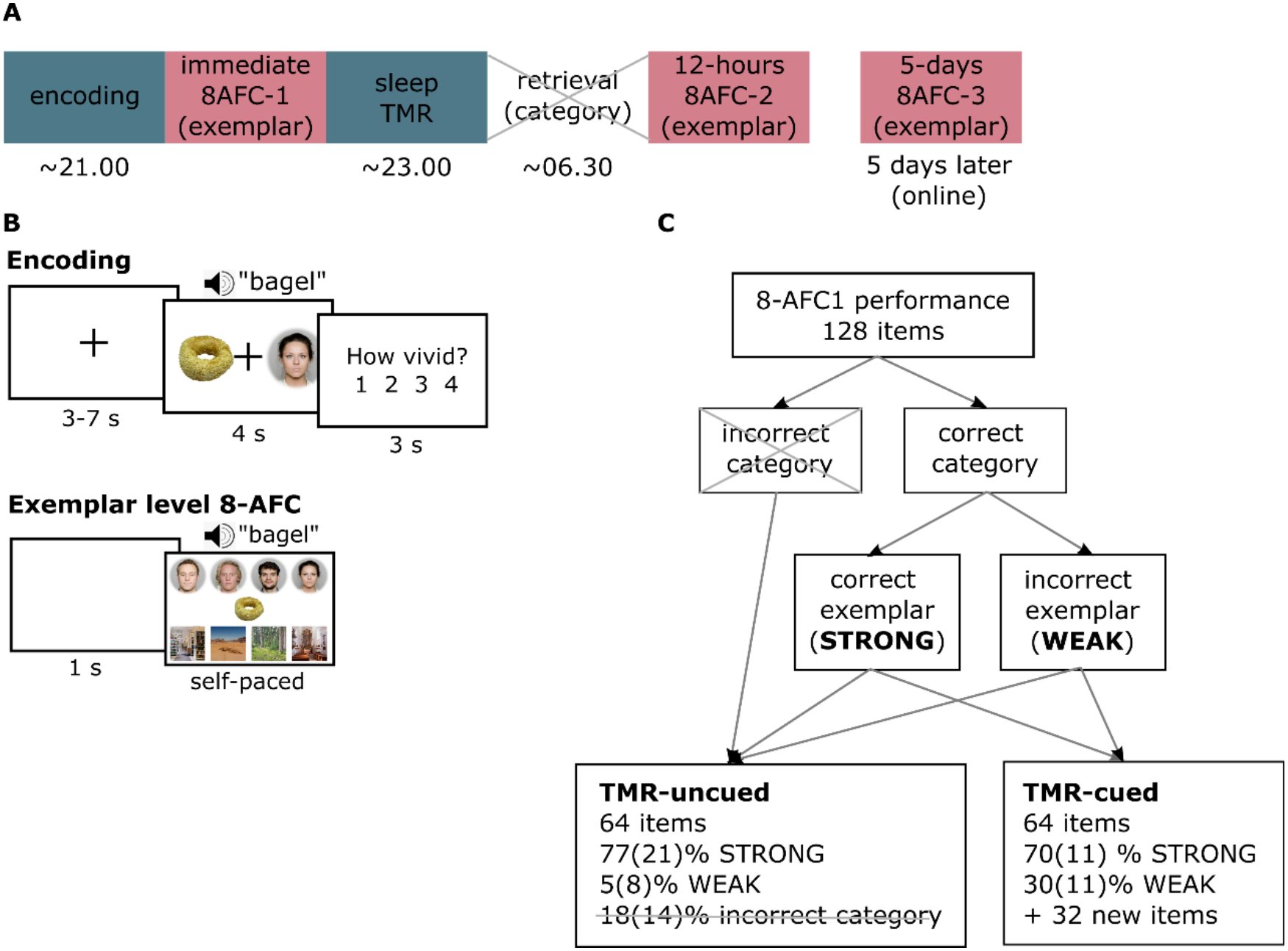
(A) Procedure and timeline of the experiment and the five-days online follow-up. The blue boxes (encoding, sleep) indicate phases of the experiment from which EEG data was analysed, and the red boxes (8AFC’s) indicate phases of the experiment that were used for behavioural performance calculation. (B) Descriptions and timelines of each behavioural experiment phase used in the analysis. (C) Diagram to indicate which items were prioritized for nocturnal cueing, based on the pre-sleep 8AFC-1 performance. The percentages in the lowest boxes indicate average proportions across participants, with standard deviations in the brackets. In (A) and (B), the crossed out elements indicate those that were not used in the analysis. The face-stimuli were used with permission from the Oslo face database^46,47^, the item-stimuli were drawn from the Bank of Standardized stimuli (CC BY)^48,49^, and the place-stimuli from Pixabay (Pixabay Content License) and Google searches, labelled for reuse with modification. The face images presented here are not identical to those used in the task but includes a selection of images in which the individuals have consented to their faces being publicly displayed.

### Memory tasks

#### Encoding

The participants performed an associative memory task in the evening (Figure 1B), in which they had to learn 128 item associations with either faces or places. Half of the associations were paired with faces, and the other half with places, and there were altogether four specific faces and four specific places. The items were presented visually on the screen for 4 s, and additionally, the word for the item (in Norwegian) was repeated auditorily from the speakers three times each. Participants were instructed to imagine an association between each item and the presented face or place, for example the person’s facial expression while using the object, or the object placed at the location. After each trial, they rated the vividness of the created association on a scale of 1–4, that is, how clearly and in how much detail they had managed to imagine the object and the face or place together (1 = not at all vivid, 4 = very vivid, clear and detailed). The response window was set to 3 s after stimulus offset. Each participant was assigned a unique combination of item-face/place pairs. The trials were divided into two runs with approximately eight minutes each and there were brief pauses between the runs. The face-stimuli were drawn from the Oslo face database^46,47^, the item-stimuli from the Bank of Standardized stimuli^48,49^, and the place-stimuli from Pixabay and Google searches, labelled for reuse with modification. All memory tasks were implemented in MATLAB R2019a, Psychtoolbox 3.0.16, and auditory stimuli were presented with a single Genelec 8020d speaker.

#### Exemplar-level 8-alternative forced choice task (8AFC)

The learned associations were tested at the exemplar level at three time points (Figure 1A-B: directly after encoding (8AFC-1), next morning after the sleep and TMR procedure (8AFC-2), and five days later online (8AFC-3). This task was self-paced, in which participants were shown each of the learned items with all faces and places used in the task, and the participant had to select, which specific face or place exemplar the item had been paired with during encoding. Just before the 8AFC-2, there was an additional category-level free association retrieval task, which was not used in the current analysis.

### TMR procedure

From the encoded items, 50% (64 items: 32 faces, 32 places) were selected for nocturnal cueing. In addition, 32 new items were introduced, that the participants had not heard before. The cueing consisted of a single repetition of each word, played from the speakers placed at approximately 50 cm distance from the participants’ heads. The words were played with jittered inter-trial interval of 4-7 s, the order of words in the full cycle of words was random, and this cycle was repeated up to 12 times during the night (totalling 1152 trials per participant). If any transitions to wakefulness, NREM stage 1, or REM sleep was detected, the cueing was paused and only continued if the participant entered stable NREM stage 2 or 3 again.

The sound level of the cues was determined on a short sound test before sleep. Here, participants were played a sequence of five (previously not heard) words repeatedly over three repetitions (15 words in total), and after each presentation, they had to indicate whether they had heard the item and understood the semantic meaning of the word. If they answered “yes”, the sound level was decreased and vice versa for a“no” answer. The final stimulation volume was calculated by taking the mean of the 3 lowest volume values with “yes” answers and multiplying it by 1.2 (i.e. 120% of the threshold). This procedure typically resulted in a level of ≈40 dB, as measured from the pillow. If participants woke repeatedly from NREM-2/3 sleep at the onset of cueing, the sound level could be reduced manually to prevent further awakenings.

After the sound test, participants were requested to lay still in their beds with eyes open for a 12-min resting state recording. After this, they heard a gentle bell from the speaker, which indicated that they were allowed to close their eyes and fall asleep.

The items were selected for cueing pseudo-randomly, based on the exemplar-level test right after encoding (8AFC-1; Figure 1C). First, only items that were correctly associated with any face or place were considered (correct category), on the assumption that if no memory trace was created during encoding, this would not be further consolidated during sleep. The final proportion of cued items was fixed so that up to 60% of items were selected among the pool that was correctly remembered to be paired with a specific face or place exemplar (parametrized as “strong memories”), and the remaining 40% had been associated with the correct category, but incorrect face or place exemplar (“weak memories”). However, due to the relatively high performance rates at 8AFC-1, the empirical proportions ended up at 70 and 30% for strong and weak memories, respectively. In addition, as weak memories were prioritized for cueing, the items that remained for the “uncued” condition were mostly classified as strong memories (Figure 1C).

The TMR presentation was implemented in MATLAB R2022a and Psychtoolbox 3.0.18. Auditory stimuli were presented with a single Genelec 8020d speaker.

### EEG acquisition

EEG was measured using 32-channel caps specialized for sleep research in the MRI scanner (BrainCap MR Sleep) and amplified with BrainAmp MR amplifier (Brain Products GmbH, Germany). Specifically, the flat Ag/AgCl electrodes were fastened to the cap, electrode FCz was set as online reference, two electrodes were reserved for EOG, three for chin EMG and one as ECG electrode, leaving 25 EEG electrodes after re-referencing against average mastoids. Data was quantified at 250 Hz sampling rate. Thick long-lasting electrode paste was used to achieve electrode impedances <5 kOhm before sleep. EEG was recorded at all phases of the memory task, except for the 5 days online test, and continuously during the full night of sleep.

### EEG preprocessing

Preprocessing was implemented with EEGLAB toolbox^50^ (v.2020_0, MATLAB 2023a), First, data from both encoding runs was concatenated. Next, sleep data was inspected manually to identify channels with unstable signal, and these were interpolated. EEG channels were then re-referenced to the average mastoids, and two chin EMG channels were referenced against the third chin EMG electrode. EEG data was filtered with a zero-phase non-causal filter at 0.1 – 35 Hz (6 dB cut-offs at 0.05 Hz and 39 Hz for high- and low-pass, respectively). Data was epoched and time-locked to all stimulus onsets with epoch durations of -1.5 – 5 s, and baseline corrected to the average across the -200 – 0 ms of the pre-stimulus signal. First, automatic probabilistic detection and rejection of trials marked as artefacts was applied. However, as the data was often contaminated with slow-wave large-amplitude artefact from sweating, additional manual inspection and trial rejection was implemented. The encoding data was further subjected to independent component analysis, and components that were automatically labelled by the *iclabel* plugin as eye (threshold = 0.7) or muscle artefacts (threshold = 0.9) were deleted before the backprojection of the components into the electrode space.

### Sleep scoring

To facilitate online sleep scoring for correct TMR administration during data acquisition, an online processing pipeline was implemented for visualization, which included a band-pass filter at 0.1 - 35 Hz and re-referencing that selected frontal, central, and occipital electrodes against their respective contralateral mastoids. Continuous filtered EEG data was additionally sleep-scored offline using the automatic algorithm YASA^51^ (v. 0.6.5, Python v.3.11.5). YASA outputs the assigned sleep stage label to the data in steps of 30 s, as well as the confidence rating for the segment belonging to any sleep stage. Sleep architecture (duration, wake after sleep onset, and sleep efficiency) and sleep-stage confidence levels during auditory cues were quantified to validate sleep and successful TMR application. One participant was discarded from sleep architecture analysis, as the first half of the night-EEG was not recorded due to technical reasons.

### Time-frequency analysis

The epoched data from encoding and sleep were decomposed using Morlet wavelet analysis (EEGLAB v. 2025.0.0., MATLAB 2023a). Time–frequency estimates were retained at the single-trial level without initial baseline correction. Analyses covered frequencies from 1 to 20 Hz, with the number of wavelet cycles increasing linearly from 1 cycle at 1 Hz to 10 cycles at 20 Hz. Power was calculated as 10*log10(real^2^+imag^2^).

To define analysis windows, time–frequency data were first averaged across all trials, separately for encoding and sleep. These average responses were normalized by division by the frequency-specific mean power in the pre-stimulus interval from -800 to -500 ms. This baseline window was selected to avoid contamination of the baseline by stimulus-evoked slow-wave activity. Electrodes and frequency ranges were specified *a priori* based on typical encoding- and TMR-related EEG responses. The sigma band was divided into high and low sigma, reflecting the supposed physiological and functional dissociation between fast and slow spindles, respectively^52^. Time windows were then defined as periods during which the trial- and group-averaged response within each channel–frequency range exceeded the 95^th^ percentile within the 0-4 s post-stimulus interval (Table 1).

**Table 1.** Channel-time-frequency window selected for statistical analysis. . Frequency bands and electrodes were selected a-priori, whereas the time-window reflects the strongest response-period (>95^th^ percentile) in the trial- and group-averaged state-specific response.

|  | Frequency (Hz) | Time (ms) | Channels |
|---|---|---|---|
| **Encoding** |  |  |  |
| Delta | 1-4 | 140-492 | Fz, FCz, Cz, F3, F4, C3, C4 |
| Theta | 4-8 | 84-328 | Fz, FCz, Cz, F3, F4, C3, C4 |
| **Sleep** |  |  |  |
| Delta | 1-4 | 516-952 | Fz, FCz, Cz, F3, F4, C3, C4 |
| Theta | 4-8 | 356-736 | Fz, FCz, Cz, F3, F4, C3, C4 |
| Low sigma | 9-12 | 1084-1708 | Fz, FCz, Cz, F3, F4, C3, C4 |
| High sigma | 12-16 | 1032-1356 | Cz, Pz, C3, C4, P3, P4 |

For item-level EEG analyses, sleep data were restricted to trials delivered successfully during N2 or N3 sleep. Baseline normalization was then performed at the single-trial level, in which the post-stimulus activity was divided by state-specific reference distributions. For encoding trials, each trial was normalized relative to the mean of all concatenated encoding baseline segments (-800 to -500 ms before stimulus onsets). For sleep trials, separate baseline distributions were created by concatenating baseline segments (-800 to -500 ms before stimulus onsets) from all trials delivered in N2 and N3, respectively. Each trial was then normalized relative to the distribution corresponding to its sleep stage. This approach bypasses the positive bias associated with normalizing each trial by its own pre-stimulus baseline^53–55^. State-specific normalization was additionally necessary because the overall EEG power spectrum differs substantially between wakefulness, N2 sleep, and N3 sleep.

For each trial, normalized power was averaged across the predefined channels, frequencies, and time windows shown in Table 1. Note that as each item was presented once during encoding, the power estimates represent single-trial responses relative to the overall wake-encoding power spectrum. However, during sleep, each item could be cued up to 12 times. Sleep responses were therefore averaged across all available cue repetitions for each item, yielding an average induced response for each item relative to the sleep power spectrum.

### Statistical analysis

Statistical analyses were conducted at the item level using generalized mixed-effects models fitted in R (version 4.2.1) with the *lme4* package^56^. Item-level models allowed us to follow individual items across time, jointly model encoding- and sleep-related predictors, and retain observations despite unequal numbers of subsequently remembered and forgotten items.

Unless stated otherwise, all models included age, sex, and item category (face versus place) as covariates, with random intercepts for items nested within participants. Effect sizes were estimated with the *marginaleffects* package^57^. For continuous predictors, we report average marginal effects (AME), representing the average change in predicted outcome probability associated with a one-unit increase in the predictor. For categorical predictors, we report average differences in predicted probability between categories. Unless stated otherwise, categorical predictors were contrast-coded, so that the reported results reflect average effect sizes across the levels of all other categorical predictors. For the EEG analysis, trials in which power in any of the analysed frequency bands deviated by more than 5 SD from the participant’s mean were excluded as outliers (0-0.1% of all trials).

#### (1) TMR effects on memory

To test whether TMR improved memory, exemplar-level correctness at post-sleep tests was modelled as a binary outcome. Items that were category-incorrect at the pre-sleep 8AFC-1 test were excluded because they were not eligible for cueing. As predictors, the model included encoding vividness (subjective ratings from 1–4), TMR-condition (cued versus uncued), test time (12-h versus 5-d) and pre-sleep baseline memory performance, coded as strong (category and exemplar correct) or weak (category correct but exemplar incorrect). Additionally, TMR-condition and test-time, as well as TMR-condition and pre-sleep performance interactions were included. This primary model was compared with models additionally including either the TMR condition, test-time, and vividness interactions, or the TMR-condition, test-time, and item-category interactions.

As a control analysis, all models predicting memory performance were repeated after restricting the data to items that elicited delta responses above each participant’s median. Cue-evoked delta activity may provide an indirect marker of auditory-evoked K-complexes, which have been associated with auditory processing during sleep^58^. This within-participant median-split approach therefore focused on items that elicited relatively strong responses, and thus may have been more likely to undergo higher-order memory processing during sleep.

#### (2) Does auditory cueing during NREM sleep elicits memory-related neural responses?

To test whether the sleeping brain differentially processed learned and novel sounds, we fitted separate linear mixed models for cue-evoked power in each sleep frequency band (delta, theta, low sigma, high sigma). In each model, cue-evoked power was predicted by TMR cue type (learned = 1, novel = 0), with a random intercept for participants. To make effects comparable across frequency bands, we report standardized coefficients (β), expressing the difference between learned and novel cues in standard deviations of the respective band’s power. P-values were Holm-corrected for the four models.

#### (3) Do the power of cue-evoked oscillatory responses during sleep relate to oscillatory power during associative encoding?

To test whether item-specific oscillatory responses during wakeful encoding were associated with subsequent responses to the same memory cues during sleep, mean power was extracted from each predefined time–frequency window (Table 1) from item-specific maps. Bivariate correlation matrices were calculated between all encoding and sleep frequency bands. Item-level correlations were computed on values z-scored within participant, thereby testing whether, relative to a participant’s own average, items eliciting stronger encoding-related responses also elicited stronger cue-evoked sleep responses while removing between-participant differences.

As a complementary analysis, analogous correlation matrices were calculated using participant-level, trial-averaged responses. These correlations assessed between-participant associations, for example, whether participants showing stronger theta responses during encoding also showed stronger theta responses to learned cues during sleep.

To test whether encoding- and sleep-related oscillations had different spatial scalp distributions and thus potentially distinct neural generators, we conducted nonparametric topographic analyses of variance (TANOVA) for dependent samples. TANOVA quantifies global map dissimilarity, which is insensitive to overall response amplitude and instead tests differences in the spatial configuration of activity across electrodes^59,60^. This analysis was implemented using freely available toolbox^61^ and FieldTrip^62^ in MATLAB R2026a.

Analyses comparing wakeful encoding and sleep topographies were restricted to delta and theta bands, which showed reliable cue-induced activity during both wakeful encoding and sleep TMR. For each participant and comparison, power within each relevant time–frequency window was averaged across trials. Each resulting scalp map was normalized by its global field power (i.e. electrode values were mean-centered and divided by their spatial standard deviation). The observed test statistic was the difference between grand-average normalized maps, equivalent to global map dissimilarity. Statistical significance was evaluated using 10 000 within-participant sleep-wake label permutations, and global map dissimilarity was recalculated for each permutation to form an empirical null distribution. The p-value was the proportion of permuted statistics greater than or equal to the observed statistic.

#### (4) Do oscillatory power during encoding and sleep predict memory performance?

To test whether encoding-related oscillations predicted later memory, we fitted an encoding-only model including delta power, theta power, and subjective vividness as predictors of item accuracy. Test time was represented by two contrasts: pre-sleep versus post-sleep performance and 12-h versus 5-d post-sleep performance. We compared this model with models including interactions of vividness with encoding delta and theta power, and interactions of test time with encoding delta and theta power.

To test whether cue-evoked sleep oscillations predicted later memory, we first fitted a sleep-only model including sleep delta, theta, low-sigma, and high-sigma power as simultaneous predictors of item accuracy, in addition to the test-time (12-h versus 5-d). We next expanded this model by adding encoding vividness, pre-sleep memory status, and encoding delta and theta power, to account for behavioural and encoding-related determinants of memory.

We additionally tested whether associations between sleep EEG power and memory varied with encoding vividness or test-time by adding the corresponding interaction terms. To elaborate on a behavioural finding of different TMR effects on strong and weak memories, we directly tested whether cue-evoked power in each sleep frequency band differed between strong and weak items.

Because encoding vividness robustly predicted later memory, we conducted exploratory analyses of its associations with encoding- and sleep-related oscillations. First, we fitted a linear mixed-effects model predicting vividness ratings from encoding delta and theta power, with age and sex as covariates and a random intercept for participant. We then tested whether encoding vividness predicted subsequent cue-evoked sleep power by fitting models with sleep-band power as the outcome and vividness as the predictor.

## Results

### Sleep and TMR validation

The average total recoding time was 6.87 h (SD = 0.37), of which total sleep duration was 5.8 h (SD = 0.70). Wake after sleep onset was 51 min (SD = 33), and sleep efficiency was 84% (SD = 10). Participants spent, on average, 175 min (SD = 29) in stage N2, and 85 min (SD = 30) in stage N3 sleep. Offline automatic sleep staging confirmed that 93.6% of the cues fell onto the N2 or N3 sleep stage (wakefulness = 3.06%, N1 = 0.89%, N2 = 39.9%, N3 = 53.7%, REM = 2.44%). The confidence ratings of the automatically assigned sleep stages for cues at N2/N3 stages was 0.87.

### TMR effects on memory

TMR had a negative effect on post-sleep memory performance (Figure 2A, Table 2), and this effect interacted with pre-sleep baseline performance. Specifically, the probability of correct memory response was worse for cued items, compared with uncued items, but this effect was restricted to weakly encoded items (AME(weak) = -10.20 percentage points (pp), AME(strong) = -0.14 pp). The pre-sleep baseline performance itself had a strong effect on memory performance, in which items that were remembered strongly during encoding had a stronger likelihood of being remembered after sleep. In addition, post-sleep memory performance was affected by encoding vividness (Figure 2B), indicating ≈3 pp increase in accuracy with each increasing level of vividness. There was also a main effect of test-time, indicating forgetting ≈13 pp after 5 d, compared with 12 h after encoding. Lastly, there was a main effect of category, with places having increased accuracy compared with faces. TMR did not interact with test-time, and the models with vividness or category interactions did not improve the model fit (both p’s > 0.72). The results remained the same if items that were unwantedly presented in wakefulness, N1, or REM sleep at least once (out of up to 12 repetitions) were discarded from the analysis. The results also remained qualitatively same if we limited the cued items to those that elicited a large delta response during sleep (median split by person as proxy for elicited K-complex).

**Figure 2.**
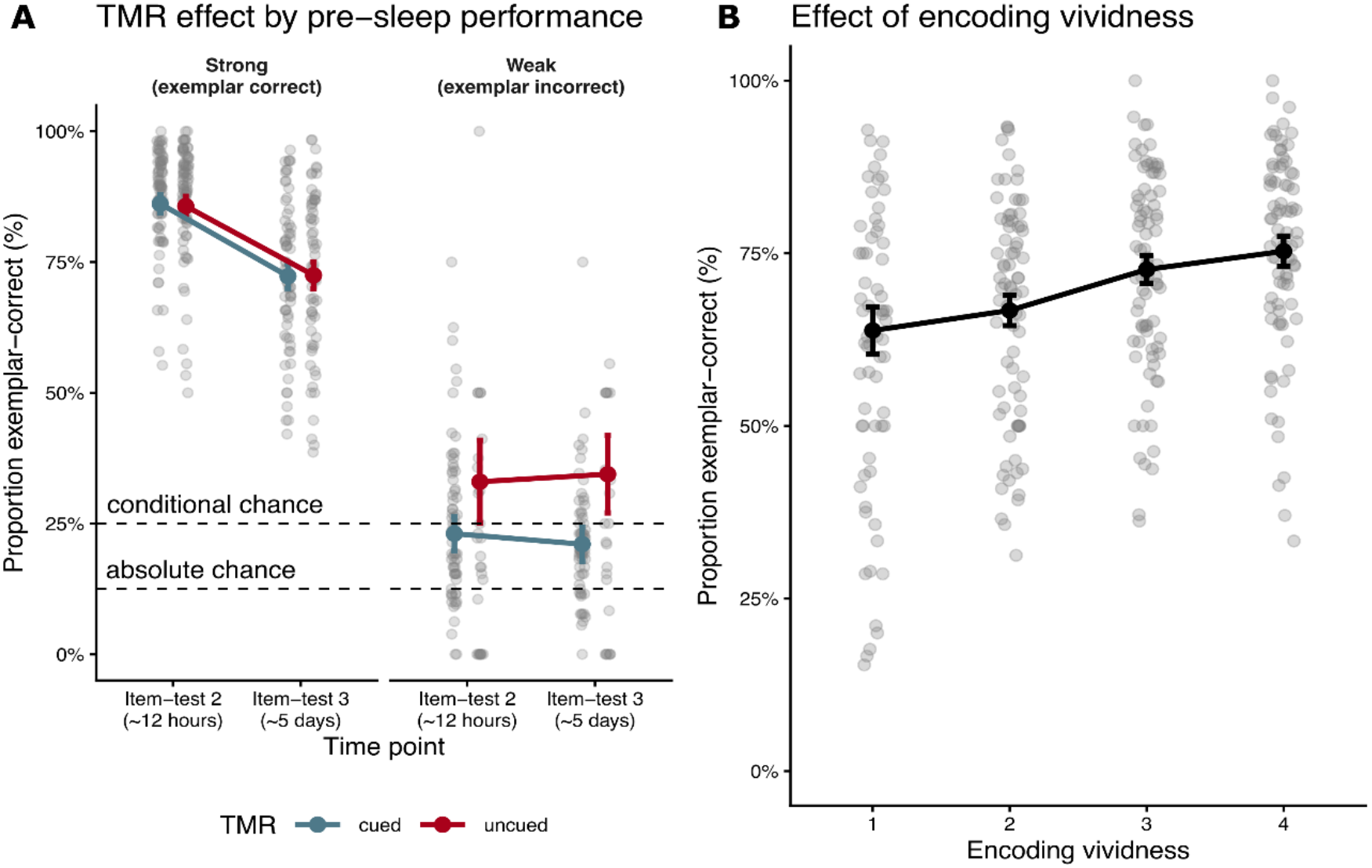
Behavioural results. **(A)** Effect of nocturnal TMR cueing on memory retention 12 hours and 5 days after encoding. Gray dots represent individual participants, and the blue and red dots indicate group estimated marginal means for cued and uncued items, respectively. Error bars indicate 95% confidence intervals. “Absolute chance” indicates the chance level across all 8 response alternatives, whereas “conditional chance” indicates the chance level for correct exemplar out of 4 faces or places, given that category was correctly remembered at pre-sleep baseline. **(B**) Effect of encoding vividness on memory retention. Gray dots represent individual participants, and the black dots indicate group estimated marginal means with error bars indicating 95% confidence intervals.

**Table 2.**
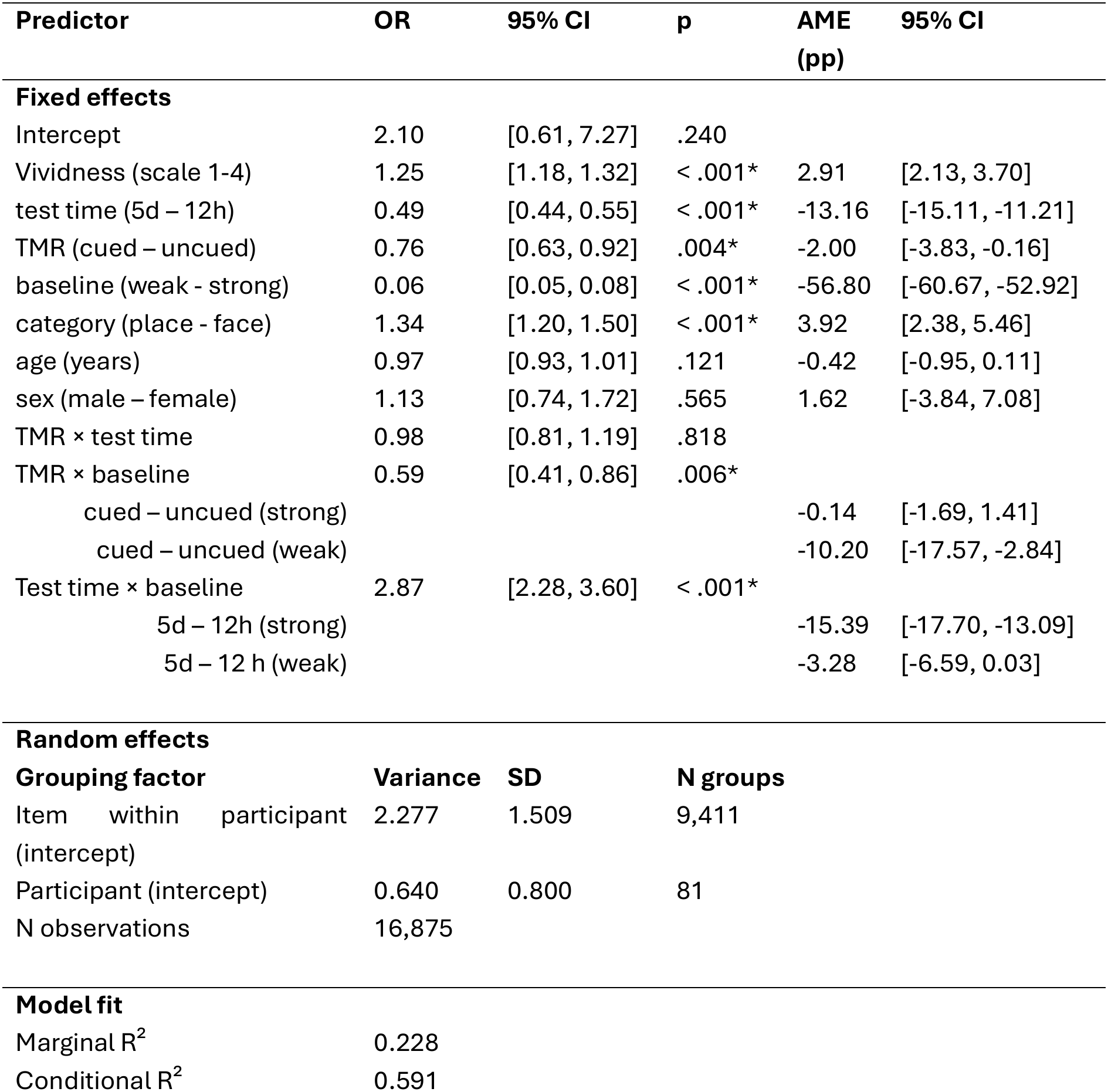
Behavioural results. Logistic generalized linear mixed model (binomial distribution, logit link). Categorical predictors were contrast-coded. OR = odds ratio with Wald 95% confidence interval; p-values are from Wald z tests. For continuous predictors, ORs are per one-unit increase; for categorical predictors, ORs compare the two levels, averaged over the levels of the other factors; for interactions, ORs are ratios of odds ratios. AME = average marginal effect on the probability of correct memory item in percentage points (pp), with 95% confidence intervals (random effects set to zero). For continuous predictors, AMEs are per one-unit increase; for categorical predictors, they are the difference between levels. AMEs for main effects are averaged over the observed values of all other predictors. For significant interactions, AMEs are given as simple effects, i.e. the effect of one factor at each level of the other (indented rows); no AMEs are given for non-significant interactions. Random-effect variances and SDs are on the logit scale.

### Sleep EEG responses to learned and novel cues

Auditory TMR cues elicited robust evoked EEG responses in delta, theta, and high sigma bands, and relatively smaller response in the low sigma band (Figure 3A, middle and right panels). We predicted induced EEG power during sleep from the categorical factor of whether an item was new or previously learned, separately for delta, theta, low sigma, and high sigma bands, and found a significant effect for the delta band (Table 3, Figure 4A). The other frequency bands did not dissociate between previously learned and new items.

**Figure 3:**
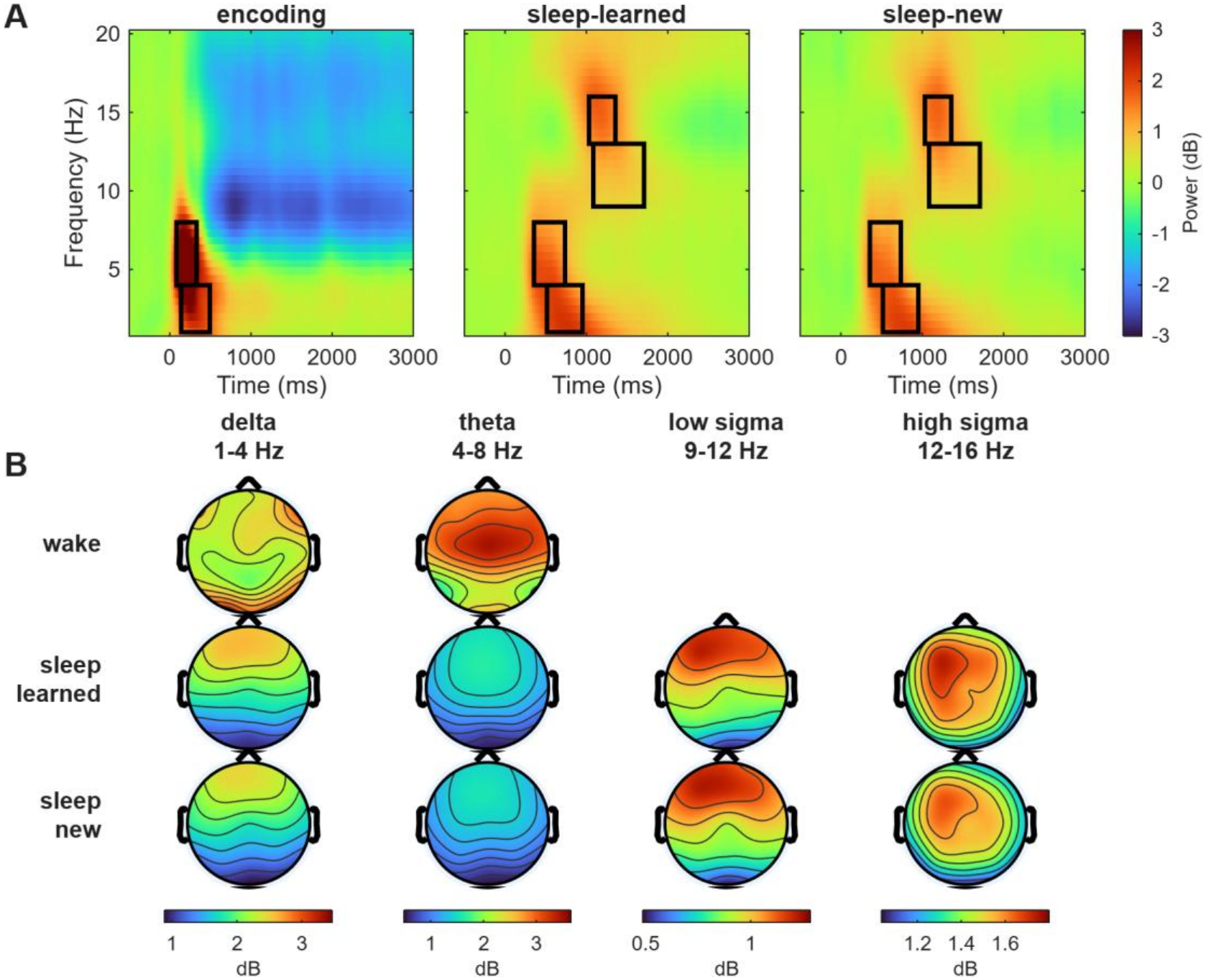
EEG time-frequency responses to memory items during wakeful encoding and sleep. **(A)** Time-frequency representations at channel Cz, averaged across all trials during encoding and sleep (separately for previously learned and new control items), time-locked to the stimulus onset. The black rectangles represent the time-and-frequency ranges selected for statistical analysis. **(B)** Topographical visualization of the power values across all EEG channels, averaged over the time-frequency windows depicted in A.

**Figure 4.**
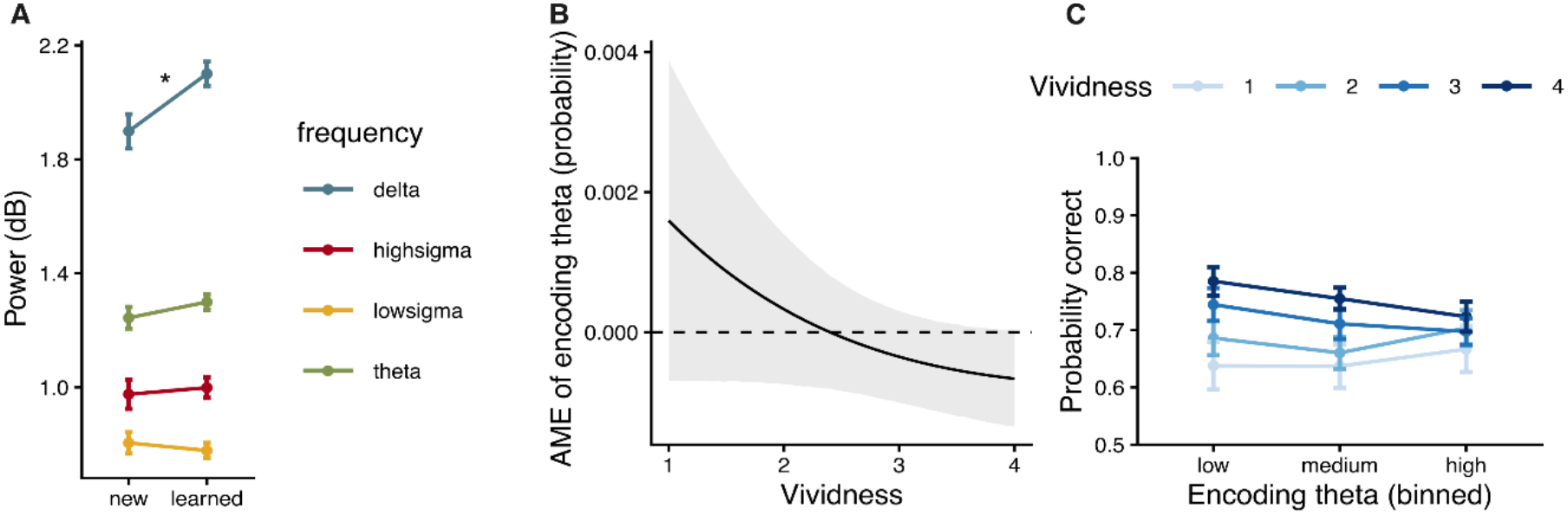
**(A)** Group averages of induced oscillatory responses, color-coded by frequency band, to new and previously learned auditory cues during sleep. The error bars represent 95% confidence intervals. *marks significant difference between new and learned items in an item-level linear regression model. **(B)** Estimated averaged marginal effects (AME) of encoding theta power at each level of vividness, with the shaded area depicting 95% confidence intervals. **(C)** Empirical data depicting the interaction of encoding theta and vividness, i.e. the average probability of a correct item at 12-h 8AFC-2 as a function of vividness and encoding theta power, binned into low, medium, and high values within each participant. The error bars represent 95% confidence intervals.

**Table 3.** Sleep-EEG results for learned vs. new items. Separate linear mixed models were fitted for each EEG frequency band (EEG feature ∼ learned + age + sex + (1 | subID)). β = difference between learned and new items in standard deviations of the respective EEG feature, with 95% confidence interval and p-values based on Satterthwaite degrees of freedom. p (Holm) = p-values corrected for multiple comparisons across the 4 models. Random-effect SDs are expressed in standard deviations of the respective feature. N = 7,665 observations from 81 participants per model.

| **Outcome** | **β** | **95% CI** | **p** | **p<br>(Holm)** | **SD<br>(participant)** | **Conditional<br>R<sup>2</sup>** |
|---|---|---|---|---|---|---|
| Delta | 0.06 | [0.02, 0.10] | .003 | .013* | 0.58 | 0.345 |
| Theta | 0.03 | [-0.01, 0.07] | .189 | .566 | 0.55 | 0.303 |
| Low sigma | -0.01 | [-0.05, 0.03] | .607 | 1.000 | 0.48 | 0.235 |
| High sigma | 0.01 | [-0.03, 0.05] | .594 | 1.000 | 0.52 | 0.265 |

### Associations between encoding- and sleep-evoked EEG responses

Memory items during encoding evoked robust EEG theta and delta responses (Figure 3A, left panel). To test whether item-specific EEG responses during encoding were associated with subsequent cue-induced responses during sleep, we calculated bivariate correlations between all wake and sleep frequency bands. As shown in Figure 5A, correlations were moderate to high among frequency bands within each state (all r’s >= 0.23, all p’s < 0.001), but no associations emerged between encoding- and sleep-induced responses (all r’s <= 0.002, all p’s >= 0.12). The complementary participant-level analysis showed the same absence of cross-state correspondence (Figure 5B), with average encoding-induced and sleep-induced responses being essentially uncorrelated across participants.

**Figure 5.**
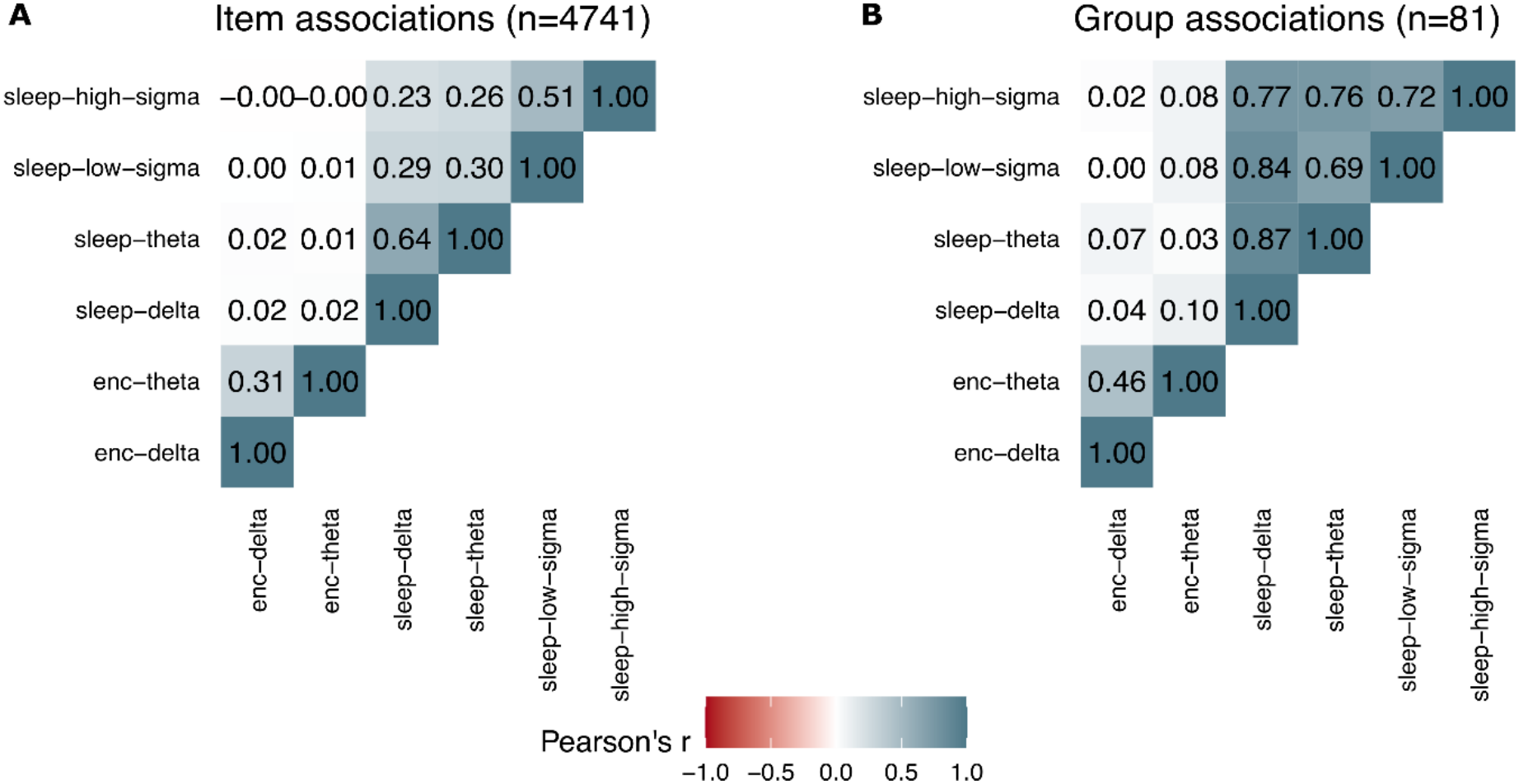
EEG associations between sleep and wakefulness. **(A)** Bivariate correlations between time-frequency responses induced during wakeful encoding and sleep across memory items, z-scored within each participant to remove individual differences. Encoding-activity represents single-trial activity, whereas sleep-items were averaged over up to 12 cue-repetitions per night. **(B)** Same associations across participants, in which activity is averaged across all trials per participant.

We next compared the scalp topographies of encoding- and sleep-induced delta and theta activity as an indirect test of whether responses in the two states might reflect different underlying neural generators. Delta activity was broadly distributed across the scalp during encoding but was maximal over central electrodes during sleep (Figure 3B). A permutation-based TANOVA confirmed that the delta topographies differed between states (global map dissimilarity = 0.88, p_perm <_ 0.001, n = 81). Theta activity was distributed across frontal and central electrodes during encoding but was more centrally confined during sleep; these topographies also differed significantly (global map dissimilarity = 0.58, p_perm <_ 0.001, n = 81).

### Oscillatory responses during encoding and sleep and memory retention

#### Encoding oscillations

We predicted retrieval accuracy from encoding delta and theta power, and a model which included interactions between delta/theta and vividness provided a better fit over the model with no interactions (Χ^2^(2) = 10.16, p = 0.006). There were no main effects of delta nor theta, but theta activity interacted with subjective vividness (Table 4). To interpret this effect, we estimated the average effect of theta at each level of vividness (Figure 4B), which revealed positive effect of theta at the lowest vividness and negative effect at the highest (translating the AME into SD units indicates that at the lowest levels of vividness, one SD increase in theta response increases the likelihood of remembering with 0.54 pp, whereas the corresponding decrease in the highest level of vividness is -0.22 pp). Alternative visualization of the average probability of accurate responses indicates that the effect of vividness was most divergent on the lowest levels of theta (Figure 4C). The additional significant effects of the model reflected those already reported in the behavioural analysis, i.e. forgetting over time, and preferential remembering of places over faces (Table 4). The interaction models with test time did not improve the model fit. Lastly, we directly predicted vividness from delta and theta activity, and found a small, significant effect of theta (b = 0.009, t(9576) = 2.32, p = 0.021, *η*^2^ = 0.0006). In sum, whereas subjective vividness was most strongly associated with later remembering, this association was moderated by encoding-related theta activity, with differences in memory probability across vividness levels diminishing at higher theta levels.

**Table 4.**
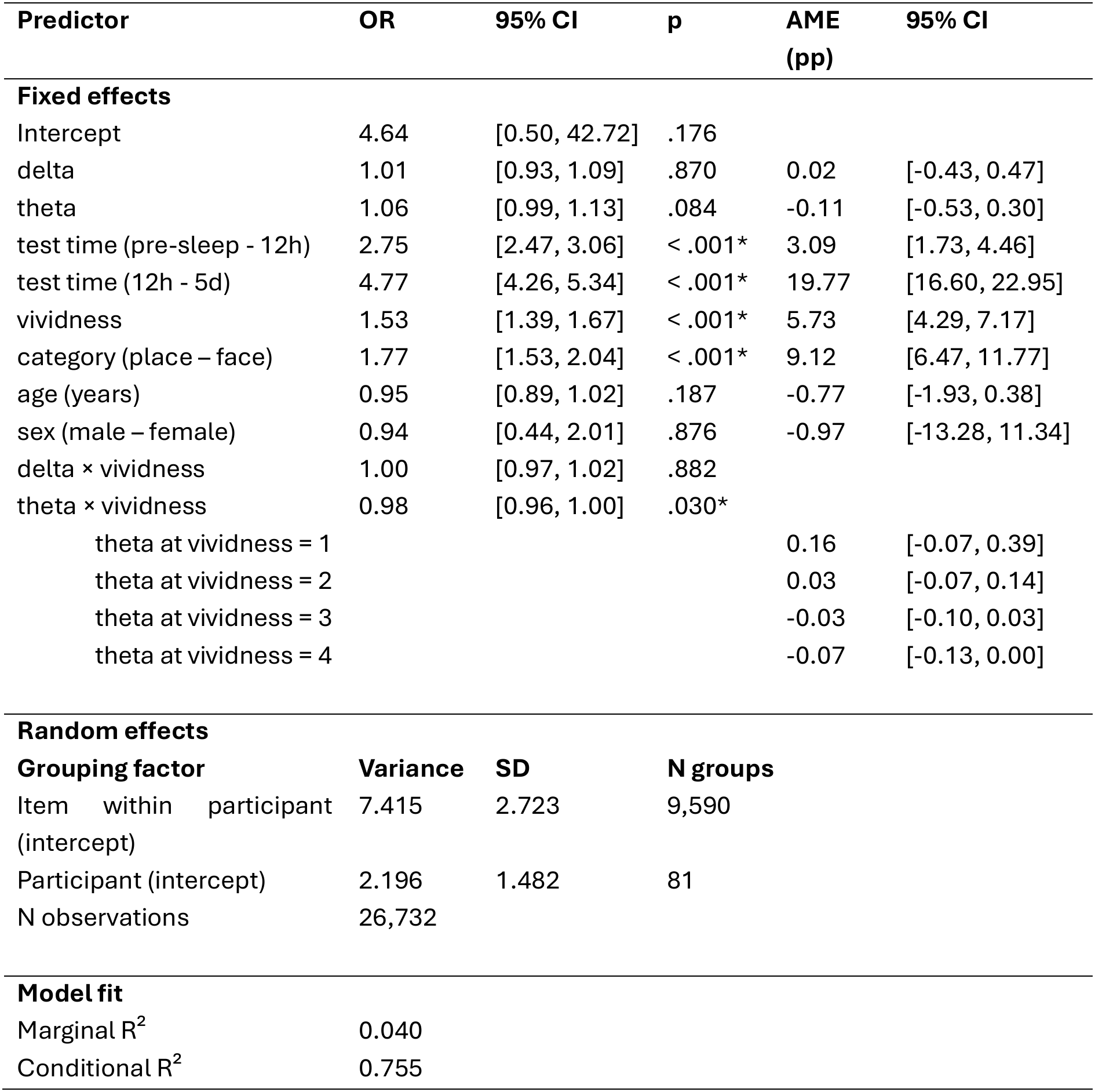
Encoding EEG results. Logistic generalized linear mixed model (binomial distribution, logit link). Categorical predictors were contrast-coded. OR = odds ratio with Wald 95% confidence interval; p-values are from Wald z tests. For continuous predictors, ORs are per one-unit increase; for categorical predictors, ORs compare the two levels, averaged over the levels of the other factors; for interactions, ORs are ratios of odds ratios. AME = average marginal effect on the probability of correct memory item in percentage points (pp), with 95% confidence intervals (random effects set to zero). For significant interactions, AMEs are given as simple effects, i.e. the effect of one factor at each level of the other (indented rows); no AMEs are given for non-significant interactions. Random-effect variances and SDs are on the logit scale.

| **Predictor** | **OR** | **95% CI** | **p** | **AME (pp)** | **95% CI** |
|---|---|---|---|---|---|
| **Fixed effects** |  |  |  |  |  |
| Intercept | 4.64 | [0.50, 42.72] | .176 |  |  |
| delta | 1.01 | [0.93, 1.09] | .870 | 0.02 | [-0.43, 0.47] |
| theta | 1.06 | [0.99, 1.13] | .084 | -0.11 | [-0.53, 0.30] |
| test time (pre-sleep - 12h) | 2.75 | [2.47, 3.06] | < .001* | 3.09 | [1.73, 4.46] |
| test time (12h - 5d) | 4.77 | [4.26, 5.34] | < .001* | 19.77 | [16.60, 22.95] |
| vividness | 1.53 | [1.39, 1.67] | < .001* | 5.73 | [4.29, 7.17] |
| category (place – face) | 1.77 | [1.53, 2.04] | < .001* | 9.12 | [6.47, 11.77] |
| age (years) | 0.95 | [0.89, 1.02] | .187 | -0.77 | [-1.93, 0.38] |
| sex (male – female) | 0.94 | [0.44, 2.01] | .876 | -0.97 | [-13.28, 11.34] |
| delta × vividness | 1.00 | [0.97, 1.02] | .882 |  |  |
| theta × vividness | 0.98 | [0.96, 1.00] | .030* |  |  |
| theta at vividness = 1 |  |  |  | 0.16 | [-0.07, 0.39] |
| theta at vividness = 2 |  |  |  | 0.03 | [-0.07, 0.14] |
| theta at vividness = 3 |  |  |  | -0.03 | [-0.10, 0.03] |
| theta at vividness = 4 |  |  |  | -0.07 | [-0.13, 0.00] |
| **Random effects** |  |  |  |  |  |
| **Grouping factor** | **Variance** | **SD** | **N groups** |  |  |
| Item within participant (intercept) | 7.415 | 2.723 | 9,590 |  |  |
| Participant (intercept) | 2.196 | 1.482 | 81 |  |  |
| N observations | 26,732 |  |  |  |  |
| **Model fit** |  |  |  |  |  |
| Marginal R<sup>2</sup> | 0.040 |  |  |  |  |
| Conditional R<sup>2</sup> | 0.755 |  |  |  |  |

#### Sleep oscillations

Next, we predicted post-sleep retrieval accuracy from TMR-induced responses in delta, theta, low sigma, and high sigma frequency bands during sleep. In the simplest model, we only had sleep EEG responses as main predictors. None of these were significant predictors of accuracy (all p’s >= 0.263). We then expanded these models to include the behavioural (vividness, condition, pre-sleep performance) and encoding-EEG variables (theta and delta responses during encoding). We also tested models where vividness or time-point interactions with the sleep-EEG predictors were included. None of these alternative models revealed any significant effects of the sleep-EEG variables. All previously described effects, including the effect of encoding theta activity and its interactions with vividness, remained significant in these models. Lastly, inspired by the behavioural results in which cueing negatively affected weakly encoded items, we directly tested the TMR-induced power between strongly and weakly encoded items separately in each frequency band, but found no significant differences in any of the bands. In sum, TMR-induced activity in delta, theta, low sigma, or high sigma bands did not predict whether the item was remembered after sleep in either time-point. In addition, subjective vividness during encoding did not predict oscillatory power in any of the frequency bands during sleep. The results stayed the same if we limited the analysis to the items that elicited large delta-response during sleep (median split by person).

## Discussion

Sleep is thought to consolidate memories that were created during encoding, yet it remains unclear how encoding and sleep factors contribute to which individual memories persist. To address this, we combined item-level measures of encoding with auditory TMR during NREM sleep. First, the sleeping brain distinguished learned from novel cues, with stronger delta responses to learned sounds. Second, encoding-related theta and delta power did not predict cue-induced delta, theta or sigma activity during sleep. This supports neither the tagging account, which predicts a positive relationship, nor the compensation account, which predicts a negative one. Moreover, TMR did not have an overall benefit of memory, and rather impaired retention of initially weak memories, contrary to the compensation account. Lastly, cue-induced EEG activity during sleep did not predict retention, and memory persistence was instead predicted by encoding-stage factors. Thus, although the sleeping brain differentiated learned from novel sounds, these responses did not predict retention, and we found no evidence that encoding-related oscillatory activity makes individual memories more reactivatable during sleep.

### Oscillatory response to auditory cues during sleep

Learned sounds elicited stronger delta responses than novel control sounds, showing that the sleeping brain distinguished memory-associated cues from unfamiliar input. Previous studies have also reported differential responses to learned and control cues during NREM sleep^35,36,63^, although the timing and frequency of these differences vary. Guttesen et al. (2024) observed an early increase in theta/alpha power followed by a sigma increase for learned cues, in a time window similar to our delta effect. In contrast, Cairney et al. (2018) reported a later sigma-band difference and Liu et al. (2023) found item-specific representations distinguishing memory from control cues only in a late window, with no early differences in low-frequency power. Here, we analysed the peak response, which occurred within the first 2 s after the cue, so the low-frequency component possibly reflects the cue-evoked slow-oscillation/K-complex response. The learned-novel distinction may thus already arise at this early stage, with spindle-related differences following later.

### TMR and memory consolidation

Contrary to our expectations, and despite the evidence that learned cues were processed differently from new sounds during sleep, TMR did not improve memory performance. This null result is unlikely to reflect insufficient statistical power, as our sample size was determined a priori from effect-size estimates of closely related work^27^ and exceeded that of many previous TMR studies. Moreover, TMR reversed the expected pattern for weakly encoded memories in which uncued weak items were retained better than cued ones. Previous findings on how prior memory strength shapes the benefits of sleep and TMR are mixed. Some studies report that sleep or TMR preferentially strengthens weak or difficult memories ^41,64,65^. Others find that TMR benefits increase with pre-sleep accuracy, except for near-perfectly learned items^40^, or that only easily learned material benefits from TMR^66^. Our results add to this heterogeneous picture by showing that cueing can even be detrimental for weakly encoded memories.

Several post hoc explanations could account for this result. First, cueing may have promoted the deprioritization of weak traces, as proposed by synaptic downscaling accounts^25,67^. Second, because weak items had been associated with an incorrect face or place exemplar before sleep, cueing may have strengthened these erroneous associations rather than the intended ones. Third, unlike most previous studies, we adopted a layered operationalization of pre-sleep performance, in which post-sleep memory was assessed at the level of the specific exemplar, while baseline performance at the gist-like category level was also taken into account. Because sleep and TMR may promote abstraction, altering the quality rather than merely the strength of memories^68,69^, post-sleep differences between cued and uncued items may partly reflect baseline variation in gist memory when this level is not controlled. Although both memory-weakening^70^ and false-memory strengthening^71^ accounts are possible, the observation that the pre-sleep baseline performance had large effects on memory retention in our study supports the hierarchical control of gist- and exemplar level memories at baseline as the strongest explanation for the behavioural results.

At the neural level, TMR cues elicited robust delta-, theta-, and sigma-band responses, yet none of these predicted subsequent memory. Previous findings here are mixed. Increased theta and spindle-band power after cues for later remembered items has been reported^39,72,73^, whereas others find robust cue-evoked responses unrelated to memory^37,74,75^. Moreover, boosting theta using transcranial alternating current stimulation after cues did not improve memory^76^. Item-specific reactivation has instead been linked to later decoded activity^35,37^ and to hippocampal ripples detectable only intracranially^77,78^. This suggests that the magnitude of cue-evoked power at the scalp primarily reflects cue processing, rather than the supposed reactivation itself.

Taken together, neither our behavioural nor neural findings, nor the mixed literature, support interpreting TMR and its induced EEG response as direct markers of memory reactivation and consolidation. Meta-analyses indicate an overall positive effect of TMR, even after accounting for publication bias^27^, yet studies examining its neural mechanisms report mixed findings or identify narrow boundary conditions, such as cueing at specific phases of slow oscillations^79–83^ or relative to preceding spindles^84^. On the other hand, positive TMR effects are also commonly reported without monitoring EEG features^85–88^, making it unclear how strongly they constrain TMR efficacy. Notably, whereas spontaneous consolidation is commonly linked to slow oscillation and spindle coupling^24,89^, the dependence of TMR on cue timing^37,84^ after spindles suggests that cued and spontaneous reactivation may not engage identical mechanisms. Moreover, consolidation also occurs during wakeful rest, when slow oscillations and spindles are absent. These oscillations may therefore provide permissive conditions for consolidation, possibly driven by hippocampal-cortical activity invisible to scalp EEG, rather than index reactivation itself.

### Linking encoding and sleep oscillations

A long-standing idea in memory research holds that the way a memory is encoded determines whether it will be consolidated during sleep, for example by tagging it for preferential reactivation. If so, the neural activity that accompanies encoding should leave a trace that can be detected when the same memory is reactivated during sleep. Here, we tested this prediction at the level of individual memories, by asking whether items encoded with stronger theta or delta power elicit distinct responses in the same or other frequency bands, when cued during sleep. We found no such correspondence. In addition, theta and delta activity showed different topographies in sleep and wakefulness.

Previous support for a link between encoding and sleep in humans comes from studies relating the two across individuals or sessions. Theta activity during learning was linked to increased spindle activity in subsequent sleep^19,42^ and individuals with stronger encoding-related theta showed more precise slow-oscillation–spindle coupling during a subsequent nap^43^. Because encoding-related theta and sleep oscillations are each related to memory performance, such associations may reflect stable individual differences in cognitive ability or physiology rather than a mechanism that acts on specific memories. Our item-level design separates these possibilities. The lack of item-specific correspondence indicates that, at least for scalp-recorded power, the electrophysiological signature of encoding does not determine how a particular memory is processed during sleep.

We focused on power because if stronger encoding reflects more synchronized engagement of memory networks, the same networks should respond more strongly when the memory is reactivated. Power was therefore a natural starting point, but it is not the only place where such a link could appear, and perhaps not the most sensitive one. First, scalp EEG signals may not generalize well across states^90^. Even if shared mechanisms support memory in wakefulness and sleep, their scalp signatures may differ because the physiological context of NREM sleep differs markedly from that of wakefulness. A common process might then appear as different power changes, or even in a different frequency band. Our event-related approach reduces this concern, as cue-induced responses were measured relative to a state-specific baseline, but it cannot account for differences in the surrounding neural sources that contribute to mixed scalp signals in each state. Even within wakefulness, similar frontal-midline theta at the scalp can arise from different source configurations^91^. Second, power reflects local synchrony rather than communication between regions. A link between encoding and sleep may instead lie in hippocampal–cortical connectivity, in the timing of cortical activity relative to slow oscillations and spindles, or in the content of the reactivated representation. Each of these can be examined at the item level, for example focusing on different time-windows^63^ or using connectivity measures, item-wise slow-oscillation–spindle coupling, or multivariate decoding of encoding-related patterns during sleep^35,90,92^. By showing that item-level power does not carry this link, the present study narrows the search and provides a framework for testing alternatives.

### Encoding determinants and durable memory

Beyond pre-sleep memory strength, subjective vividness and its interaction with encoding-related theta power predicted retention 12 h and 5 d after learning. Encoding-related theta has frequently been linked to later memory^16^, and our findings show that this relationship extends to retention over several days. Theta power and vividness were associated, but their interaction indicates that the two are not interchangeable. Theta may capture activation of cortical memory networks, while vividness ratings reflect the quality of the generated mental image^93^, but as subjective judgments likely also reflect a broader appraisal of the learning experience^94^. Memory declined from the first to the fifth day, but neither encoding nor sleep-related measures interacted with time. Encoding factors thus appear to set the initial strength of a memory, which is then carried forward. However, this conclusion should be treated with some caution, because two retention intervals give limited information about the course of forgetting. Nonetheless, our results raise the possibility that processes during encoding play a larger role in determining memory durability than the neural responses we measured during subsequent sleep.

### Limitations

Beyond the generalization gap and the focus on power discussed above, several limitations concern the design and analysis. Because uncued items have no sleep EEG equivalent, we prioritized weakly encoded items for cueing to maximize the number of trials for sleep EEG analyses, expecting that a behavioural TMR benefit would still be detectable. Combined with high pre-sleep memory performance, this left fewer comparable uncued items with preserved gist but impaired exemplar memory. As this is where the only behavioural TMR effect emerged, this effect should be interpreted with caution and replicated in a design that balances cued and uncued items across pre-sleep performance levels. In addition, encoding activity was estimated from a single presentation, whereas cues were presented up to 12 times during sleep. This asymmetry contrasts initial encoding with repeated reactivation, and the noisier single-trial encoding estimates would attenuate any item-level association between encoding and sleep responses. It also remains unclear whether the number of cue repetitions was optimal, or poorly timed to affect consolidation, particularly as cues were presented without regard to slow-oscillation phase. Rating vividness during learning may itself have influenced encoding, and testing the same items one and five days after learning may have acted as retrieval practice, reducing differences in forgetting. Although trials with cue-induced arousals were excluded from the EEG analysis, cues may still have subtly disturbed sleep, and our findings from young adults studied during a single night may not generalize to other populations or sleep conditions. Finally, normalizing single-trial responses to a state-specific baseline reduced broad spectral differences between wakefulness and sleep, but it discarded the immediate pre-stimulus period. During sleep, this period reflects whether a cue arrives during a slow-oscillation up- or down-state or after a spindle, which may shape both the induced response and TMR efficacy. Because each item was cued many times and may therefore have been presented across different slow oscillation phases, offline sorting of items by cue phase was not feasible.

## Conclusion

Memory persistence over five days was predicted by how memories were encoded, including their initial strength, subjective vividness, and its interaction with encoding-related theta activity, rather than by what happened to them during sleep. The sleeping brain distinguished learned from novel sounds, yet this differential processing neither predicted later memory nor produced a behavioural benefit of TMR. Testing a long-standing assumption at the level of individual memories, we also found that encoding-related theta and delta power did not predict how a memory responded to its cue during sleep. Together, these results show that a cue-evoked sleep response is not, by itself, evidence of memory reactivation or consolidation. Distinguishing the processing of learned cues from reactivation that changes memory will be essential for understanding how sleep shapes memory, and item-level experimental procedures and analyses provide a way to make this distinction.

## Data and code availability

Individual-level data availability is restricted as participants have not consented to publicly share their data. The data may be available upon reasonable request, given appropriate ethical, data protection, and data-sharing agreements. Requests can be submitted to the principal investigator (Prof. Anders Fjell, University of Oslo). Analysis scripts are available at https://github.com/LiisaRaud/Acon_TMR_EEG.

## Acknowledgments

We thank all research assistants on the project for their assistance with participant recruitment and data collection, including the overnight sleep recordings.

During the preparation of this manuscript, the authors used Claude (Anthropic) to assist with writing and debugging analysis code, creating data visualizations, editing and restructuring text, and providing feedback on the arguments in relation to the published literature. All code was reviewed and validated by the authors, all text was reviewed and revised by the authors, and all cited references were verified by the authors. The authors take full responsibility for the content of the publication.

